# An Analytic Platform for the Rapid and Reproducible Annotation of Ventilator Waveform Data

**DOI:** 10.1101/568386

**Authors:** Gregory B. Rehm, Brooks T. Kuhn, Monica K. Lieng, Irene Cortes-Puch, Jimmy Nguyen, Edward C. Guo, Jean-Pierre Delplanque, Nicholas R. Anderson, Jason Y. Adams

## Abstract

Algorithmic classifiers are crucial components of clinical decision support (CDS) systems needed to advance healthcare delivery. Robust CDS systems must be derived and validated via creation of multi-reviewer adjudicated gold standard datasets. Manual annotation of physiologic data such as mechanical ventilator waveform data (VWD) can be time-consuming, and lacks methodological consistency in dataset development. To address these issues, we have created a system for annotating and adjudicating VWD called the Annotation PipeLine (APL) to optimize VWD annotation by expert reviewers. APL combines visual assessment of waveform characteristics with metadata display, enabling inclusion of quantitative thresholds into annotation decisions by reviewers. APL also includes specific features for resolving multi-reviewer disagreements and generating gold standard data sets. APL’s unique combination of methods and open source framework may accelerate the creation of CDS algorithms for ventilator management, and may serve as a model for future research into physiologic waveform annotation systems.

## I. Introduction

Algorithmic classifiers promise to play a central role in analysis of different types of quantitative healthcare data, such as high-volume streaming waveform data gathered from physiologic monitoring devices. By utilizing expert-derived rules and machine learning, algorithmic classifiers can support CDS systems that can function with complex data types, and patient and disease heterogeneity. High-performance model development requires well characterized datasets annotated for salient events by expert clinicians through a process of multi-reviewer adjudication.(1-3) Accurately and reproducibly annotating data requires substantial investment of effort from time-limited, expert reviewers, which to date has limited the size and generalizability of validated datasets.(4-7)

These challenges are salient in research related to mechanical ventilation (MV). Incorporating algorithmic classifiers to MV practice has the potential to detect diagnostic events, pathologic phenomena such as patient-ventilator asynchronies (PVA), and improve clinical outcomes.(8-16). To develop and validate high performance classifiers, clinicians have historically been required to manually annotate large volumes of ventilator waveform data (VWD) by visually analyzing the shape and magnitude of air flow and pressure data.(7-10, 13) However, this process is time-consuming, subjective, and has been marked by low inter-rater agreement due to visual classification criteria based on “expert opinion”, as well as data management challenges resulting from the high volume of data and the laborious manual annotation process.(7, 10, 17) Previous groups have developed waveform annotation software that can handle large volume data and output categorization results, but these efforts lack the specific functionality necessary for overcoming problems inherent with annotating VWD.(18-30). These challenges are: 1) consistent annotation workflows do not exist for VWD annotation, which has led to the creation of datasets that are potentially non-reproducible across medical centers (17), and 2) large, high-quality, multi-reviewer adjudicated datasets of VWD are highly time-consuming to generate for expert clinicians.(7)

With these challenges in mind, we sought to address gaps in the field of VWD annotation by developing the Annotation PipeLine (APL). APL supports expert annotation of ventilator data and builds on concepts from existing platforms for annotating data other than VWD.(31,32) Utilizing APL, we have been able to annotate and provide dual clinician validation for over 315,000 ventilator recorded breaths across two different datasets. These data comprise some of the largest recorded breath-level, multi-clinician adjudicated datasets for use in developing algorithms to improve the delivery of MV.(1,7,9,10,13,33) APL’s fundamental design features allow for consistent and efficient annotation of VWD, and may be broadly applied in future research to improve efficiency, reproducibility, and fidelity of medical waveform annotation and classifier development.

## II. Methods

APL is a free, publicly accessible, web-based annotation platform developed since 2015 to support studies at University of California Davis (UCD) utilizing VWD collected from mechanically ventilated patients.(34) All VWD were collected under an approved IRB protocol and do not contain protected health information. Data were collected from Puritan Bennet Model 840^TM^ (PB-840) ventilators (Covidien, Dublin, Republic of Ireland) and stored in multiple sequential files up to two hours in length.(34,35)

APL uses an open source stack including: ventMAP software for extraction of waveform metadata, the Python Flask web framework, the JavaScript library Dygraphs for graphing of VWD, Redis for storage, and Docker for APL deployment.(1, 36, 37) APL’s source code and install instructions are available on GitHub for modification or use of APL.

### A. Improving Consistency of the Annotation Process

APL was built to support objective classification of VWD by displaying clinically-relevant quantitative metadata to help improve the consistency of the annotation workflow. After data collection, VWD is uploaded to APL in CSV format, is preprocessed for graphical display, and automatically split into individual breaths. Each breath is processed to generate clinically relevant metadata pertinent to the classification of VWD using ventMAP (Figure 1).(1,38) Examples of metadata generated include the inspiratory tidal volume (the volume delivered by the ventilator with each breath) and the expiratory time (the duration of each exhalation). All derived metadata were determined to be clinically relevant by four critical care clinicians (JYA, BTK, ICP, JN). Metadata are cached in a CSV file and then retrieved by Flask to generate a breath-level drop-up table that displays breath-specific metadata and annotation options. Display of metadata in a drop-up table mimics the user interface of a mechanical ventilator and was designed to facilitate use of APL by expert reviewers (Figure 2).

**Figure 1:**
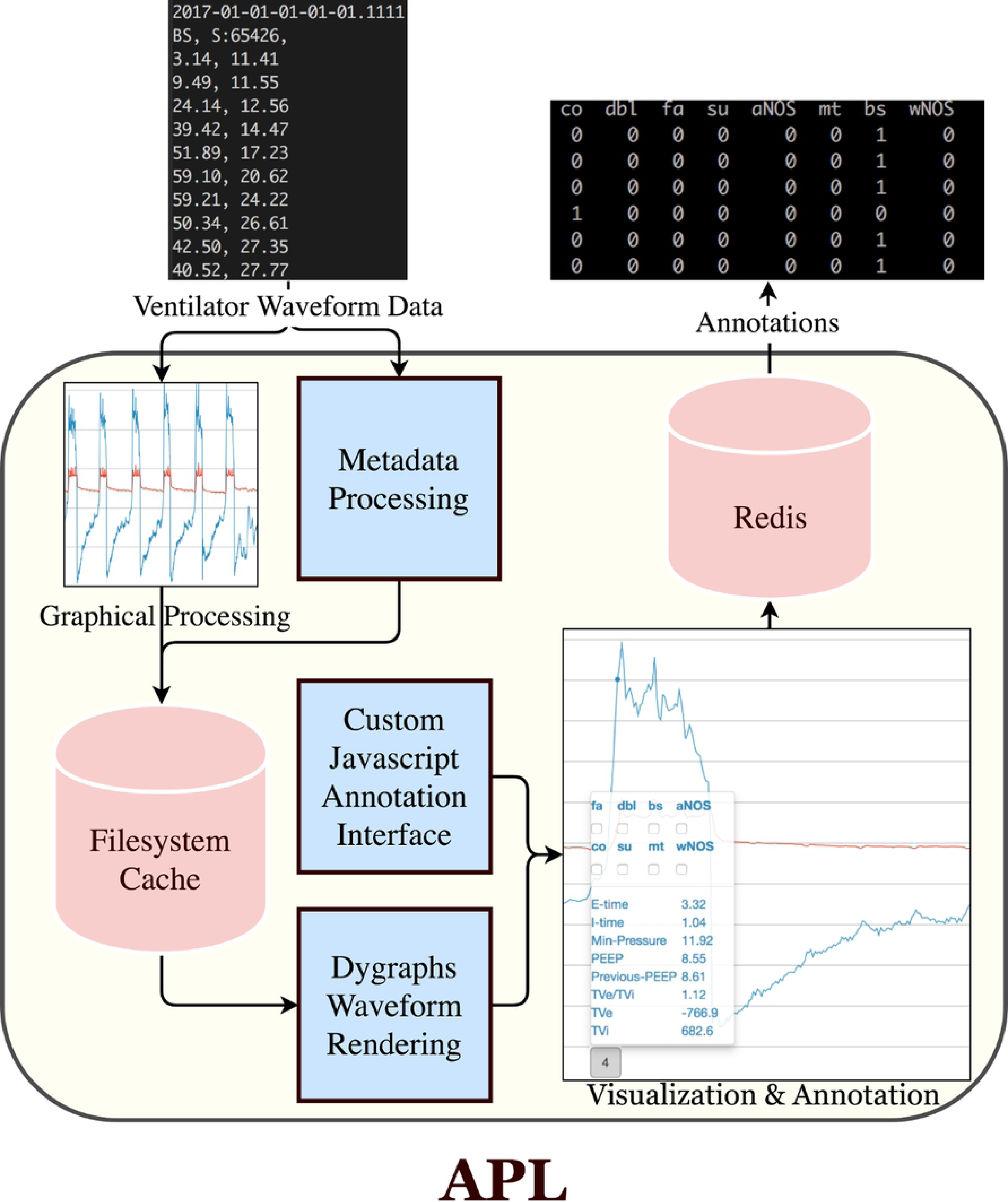
Here we display the flow of data within APL as it is processed by our software architecture. Raw VWD is uploaded to APL and then processed for graphical rendering, using ventMAP metadata processing to extract clinical metadata from each breath. Results from processing are stored on the filesystem. Next, we visualize both VWD, and perform breath-level annotation by utilizing Dygraphs with our own custom annotation overlay. All annotations are stored in Redis, and can be output by the user into CSV format.

**Figure 2:**
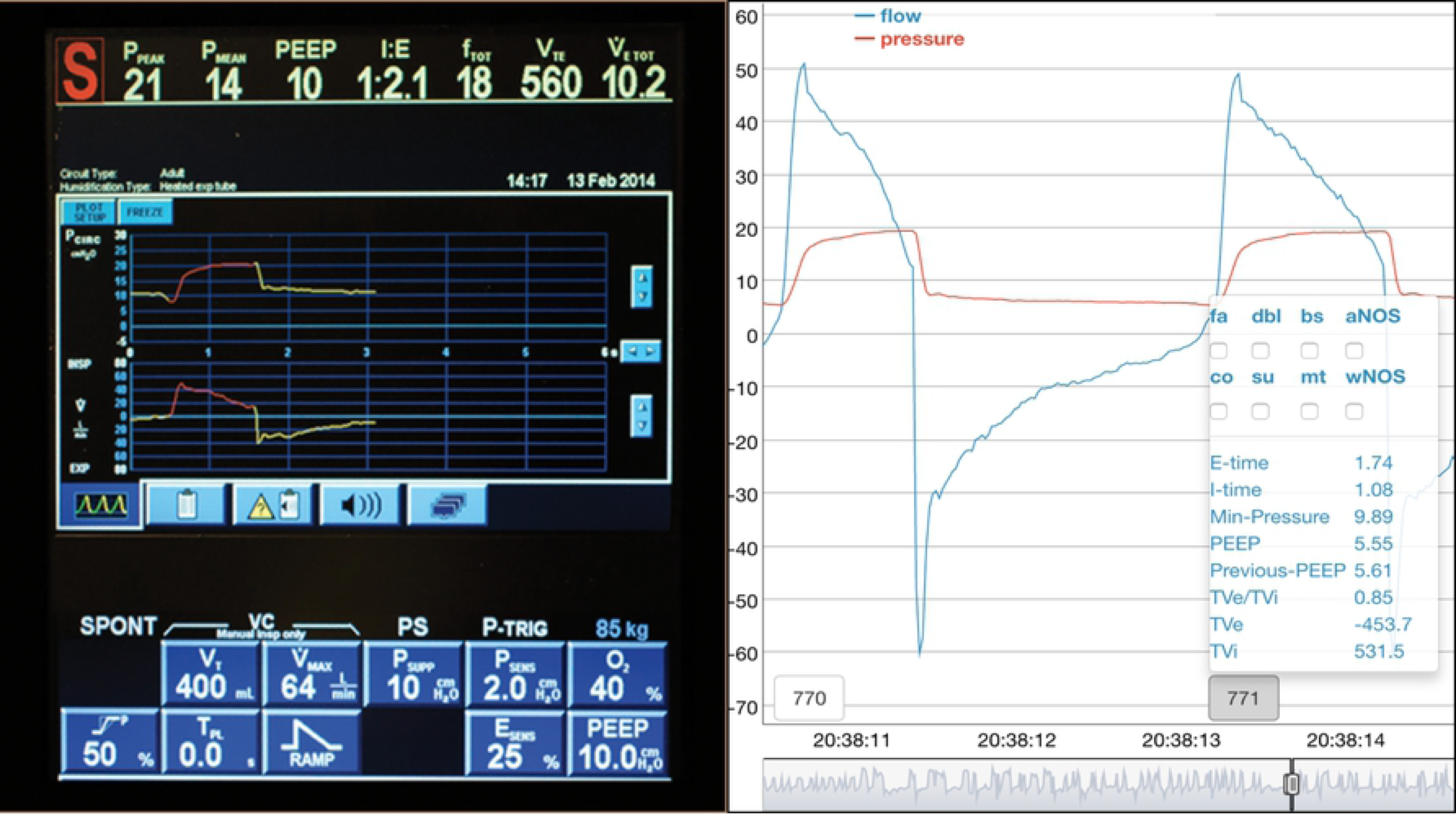
We designed APL to be familiar to clinicians, rendering information analagous to the user interface screen of a mechanical ventilator. Left, an image of the graphical user interface of a mechanical ventilator. Right, APL pressure (red) and flow (blue) waveforms with customizable breath-level metadata and point-click annotation. Breath interpretation at the bedside relies on both waveform and metadata analysis. APL presents both waveform morphology and breath-level metadata for identification of threshold-defined events.

APL also incorporates functionality to support online reconciliation of discordant annotations to improve consistency and reproducibility of the annotation process. This functionality is possible because APL is an online web platform and can store all user annotations centrally in the Redis database. Stored annotations can then be retrieved later by the Python Flask web framework to compare classification decisions by different reviewers. Differing classifications are displayed by Dygraphs as a red label that contains the cause for classification disagreement. Dygraphs displays a green checkbox above a breath if no disagreement for a breath is found.

The annotation view (AV) functionality provides the ability to select subsets of available metadata and breath classifications to display when reviewers are performing annotation. AVs are aimed at reducing information overload for the user and simplifying the annotation process by targeting it to specific project requirements. All AVs are stored in Redis, and new AVs can be created by a user by selecting the specific metadata and breath classification desired.

### B. Improving Speed of the Annotation Process

To streamline the end-to-end annotation workflow, we designed a unified system of annotation and visual display termed ‘on-screen annotation’ that allows for simultaneous display of waveform morphology, metadata, and annotation choices for every breath. (Figure 1b) On-screen annotation is accomplished by coding customized instructions in JavaScript that render drop-up tables containing metadata, annotation choices, and server callback logic. We then ensure that Dygraphs runs our overlay software after rendering of VWD. Using this overlay interface, we incorporated additional functionality to improve the speed of how annotations are performed, such as the option to perform single or multi-breath classifications.

## III. Results

Before starting the annotation process, users typically visit the Settings page to choose an AV (Figure 3). While users may choose a pre-defined AV, project leaders can create an AV, requiring that all other reviewers utilize that view to enforce consistency of annotations and outputs for a dataset. When an AV is created, leaders choose from a variety of annotation options and metadata available for display (Figure 3). When finished, the view can be named and saved to be used by other project members. In our initial work, we utilized AVs to create two different gold-standard datasets for PVA and ventilator mode (VM) classification.

**Figure 3:**
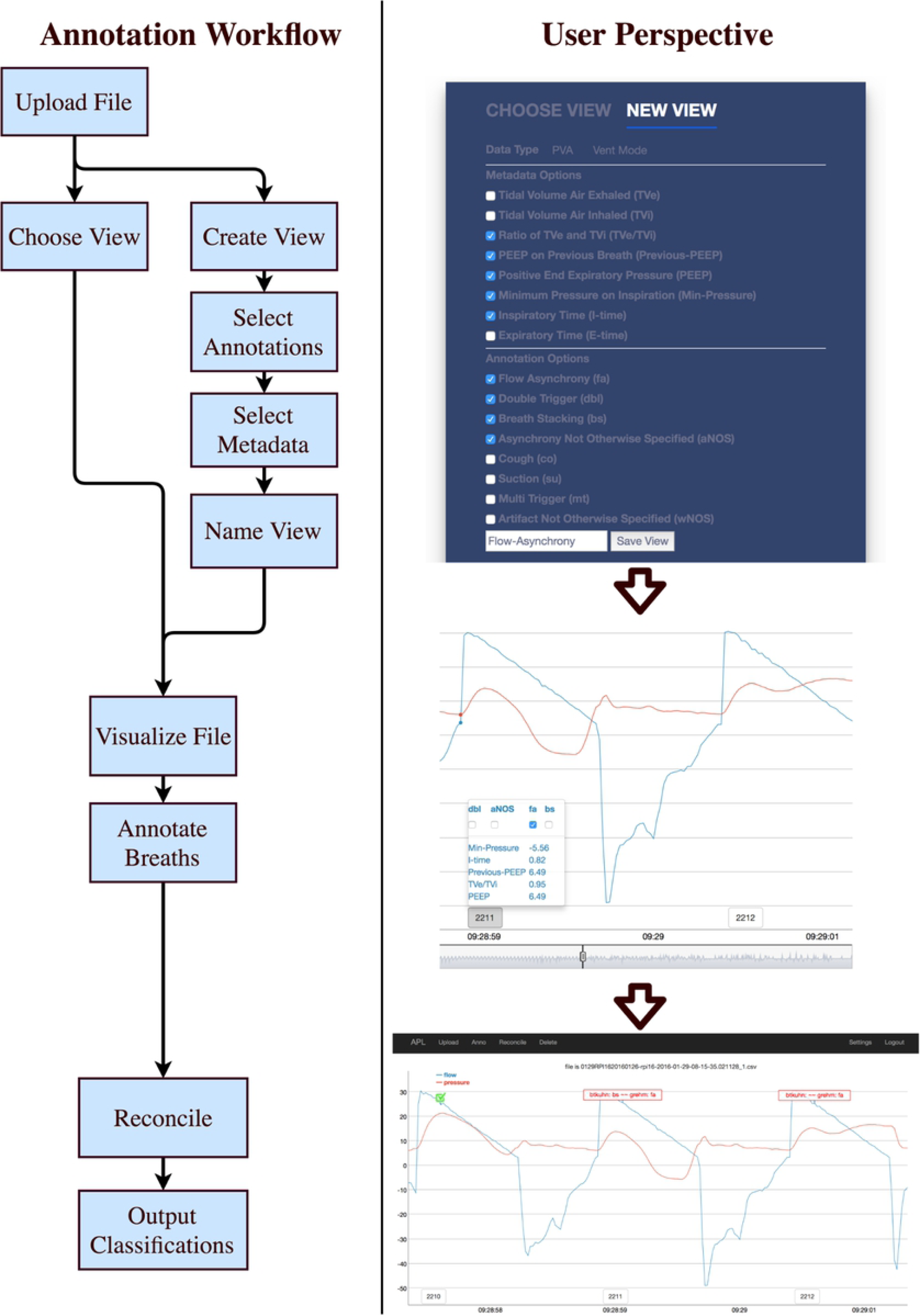
We display a typical workflow for a user when using APL, and screenshots from APL that display typical uses and functionality. On the left, annotation workflow discusses the individual steps a user would take to annotate a file. On the right three images are shown: the first showing how a user would create a new annotation view, the second displays how the annotation process occurs on ventilator waveform data. The third shows how reconciliation occurs with discordant classifications highlighted in red, displaying exactly which classifications are mismatched. Agreements are displayed as a checkbox.

Once a view is selected, uploaded VWD files can be graphically displayed for annotation. If reviewers only wish to annotate portions of a file, they can zoom in and out of specific breath regions using Dygraphs to focus on regions of interest. When a specific breath is annotated, the user can click on its breath number, and a drop-up window will appear. Drop-up windows display both quantitative metadata and annotation options allowed by the current AV (Figure 2). Users can annotate a breath by selecting the checkbox corresponding to the specific annotation desired. A user can also annotate multiple consecutive breaths using multi-breath functionality where the user annotates the first and the final breaths in a series, and all intermediate annotations will be automatically completed.

After multiple users annotate the same file, reconciliation of discordant annotations can occur. Reconciliation functionality displays VWD using an interface similar to annotation, but highlights disagreements in red and agreements with green checkboxes. Reviewers can evaluate each disagreement simultaneously and collaborate to resolve them. Once finished, users can output a CSV file containing all gold standard breath-level annotations.

Using APL, three critical care clinicians from UCD (BTK, JN, and JYA) annotated breaths for 18 different classifications from 254 VWD patient files, and generated two datasets totaling 318,676 multi-clinician adjudicated breaths. To our knowledge, this collection represents some of the largest reported datasets of breath-level, multi-clinician adjudicated VWD (Table 1).

**Table 1:** Existing multi-clinician adjudicated datasets of VWD. Number of classes corresponds with the number of annotation decisions available to the clinician as they performed annotation, including normal breaths. Exact counts for number of breaths annotated by Thille were not mentioned in the literature, so a range was extrapolated from the number of patients enrolled in their study and the respiratory rate of patients.

| **Dataset** | **Number of Annotated Breaths** | **Number of Breath Classifications** |
| --- | --- | --- |
| Pohlman 2008 (39) | 391 | 2 |
| Blanch 2012 (40) | 1,024 | 2 |
| Gholami 2018 (41) (*using ExpertLabeler*) | 1,377 | 3 |
| Sottile 2018 (7) | 2,500 | 4 |
| Colombo 2011 (15,17) | 3,731 | 4 |
| *Adams 2017 (1) (using APL)* | *9,719* | *8* |
| Thille 2006 (42) | 35,580-58,110 | 3 |
| Beitler 2016 (10) | 143,199 | 1 |
| *Ventilator Mode (using APL; in submission)* | *308,957* | *10* |

## IV. Discussion

Creating datasets of large, multi-clinician annotated VWD for MV research is limited by a lack of accessible, efficient, and reproducible multi-reviewer annotation methods. We developed the Annotation PipeLine (APL) to remedy these issues for VWD annotation and perform structured output of results. APL uses breath metadata to create hybrid visual-quantitative classifications of VWD. Incorporation of quantitative and qualitative methods into a web-based visual annotation interface may reduce reviewer subjectivity in VWD classification, and enables efficient resolution of reviewer disagreements to generate gold standard data sets to drive classifier development. Using these methods, our team has annotated 318,676 dual-adjudicated breaths for use in development of VM and PVA detection algorithms, which represents one of the largest existing collections of multi-reviewer annotated VWD.(1,7,15,17,39,40)

Historically, VWD annotation has been performed either by hand, use of electronic spreadsheets, and most recently by the web-based ExpertLabeler system.(41) A major limitation of these methods is that classifications are performed by visual criteria alone which are subject to high inter-rater bias due to reviewer subjectivity and local practice pattern.(17) APL addresses this limitation with the development of methods to support the simultaneous use of waveform morphology and metadata derived from VWD in classification decisions. To our knowledge, no MV-specific annotation package exists that is able to compute breath level metadata, display it on-screen, and allow multi-reviewer annotation and adjudication of events.

Technologically, APL builds on previous work related to annotation platform development and features novel contributions of its own. Our work to display VWD builds on previous platforms that provide annotation of generic physiologic data such as AcqKnowledge, ChronoViz, and BEDA, which have exemplified how to perform annotation on temporally archived waveform data.(18,20,43-45) APL’s combined morphologic and metadata display functionality also builds upon several previous studies that have derived higher-level metadata from raw input streams for purposes of display and annotation.(26,46-49) For example, Jing et al developed the ability to annotate multiple electroencephalogram (EEG) waveforms simultaneously.(50) Similarly, APL extends existing work developed in natural language processing applications for inter-rater reconciliation to a new domain of streaming physiologic waveform annotation.(31,32) Finally, APL contributes novel functionality including the use of drop-up windows with combined metadata and annotation capabilities, and the addition of annotation views that enable different metadata and annotation capabilities to be enforced across teams and customized to specific research needs.

Our work is limited by use of data and clinicians from a single center, and APL is presently limited to processing VWD from a single ventilator. Future work will generalize APL’s data processing capabilities to other ventilators and medical devices by exploring the use of emerging waveform encoding standards.(51) In addition, APL has only been used by four clinicians to date and further modifications may be required to meet the needs of other researchers. Changes to APL may also enable additional investigation into the reproducibility of waveform classification, and may facilitate the development of consensus guidelines for annotating VWD.(17, 52, 53) To promote further research, we have freely released our code on GitHub so that the wider research community may modify our work.

Creating effective algorithms that can detect the presence of common diseases and pathologic respiratory phenomena for CDS will require large, gold-standard datasets of VWD. In order to create generalizable datasets, researchers will also need consensus definitions for MV-related phenomena. To satisfy these needs we developed APL, the first open-source, multi-reviewer MV visualization and annotation platform that allows for the use of multi-epoch annotation to speed the classification process, customizable annotation views to constrain and standardize class assignments, reconciliation of inter-rater classification disagreements, and the simultaneous display of waveform morphology and breath level metadata to reduce classification subjectivity. While APL is specific to MV, the concepts APL utilizes can be applied to other types of high volume, physiologic waveform data. In summary, APL represents a new, effective tool to improve the consistency and speed of generating multi-reviewer gold-standard VWD datasets that will help drive the development of novel CDS systems.

